# In-vitro virucidal activity of hypothiocyanite and hypothiocyanite/lactoferrin mix against SARS-CoV-2

**DOI:** 10.1101/2020.11.17.387571

**Authors:** Luca Cegolon, Mattia Mirandola, Claudio Salaris, Maria Vittoria Salvati, Cristiano Salata, Giuseppe Mastrangelo

## Abstract

SARS-CoV-2 replicates efficiently in the upper airway during prodromal stage with resulting viral shedding into the environment from patients with active disease as well as from asymptomatic individuals. So far, virus spread has been exclusively contained by non-pharmacological interventions (social distancing, face masks, hand washing and several measures limiting business activities or movement of individuals)^1,2^. There is a need to find pharmacological interventions to mitigate the viral spread, supporting yet limiting the existing health protection measures while an effective and safe vaccine will hopefully become available. Hypothiocyanite and lactoferrin as part of the innate human immune system were shown to have a large spectrum of cidal activity against bacteria, fungi and viruses^2,3^. To test their virucidal activity against SARS-CoV-2 we conducted an in-vitro study. Here we show a dose-dependent virucidal activity of hypothiocyanite at micromolar concentrations, slightly improved by the presence of lactoferrin. The two substances are devoid of any cytotoxicity and may be administered combined by aerosol to exploit their antiviral activity at the port of entry (mouth, nasal cavity, conjunctiva) or exit (mouth, through emission of respiratory droplets) of SARS-CoV-2 in the human body. Furthermore, aerosol with hypothiocyanite and lactoferrin combined could also have a therapeutic effect in the lower respiratory tract, at the level of gas exchange units of the lung, preventing the devastating infection of alveolar type II cells where ACE2 is highly expressed. An in-vivo validation of in-vitro results is urgently required.

---

A critical problem with SARS-CoV-2 the etiological agent of COVID-19 is the high communicability of the infection, with a base reproductive number (R_0_) estimated to fall within the range of 2-3, with a mean of 3.28, a median of 2.79 and an interquartile range of 1.16, similar to SARS-CoV-1 (R_0_ = 2–5), but much higher than MERS-CoV's (R_0_ = 0.5)^4–6^. R_0_ > 1 means self-sustaining spread of the disease unless infection prevention and control (IPC) measures are enforced to curb the viral transmission^7^. Differently from SARS-CoV-1, SARS-CoV-2 seems able to replicate efficiently in the upper airways epithelium also during the prodromal stage, when large amount of viruses can be shed into the environment from asymptomatic/pre-symptomatic individuals also during the incubation period, which can stretch up to 14 days^1,2,8^. Asymptomatic/pre-symptomatic individuals make up about 90% of patients infected by SARS-CoV-2, thus rendering the spread of COVID-19 much easier than SARS, whose communicability was limited to the critical active phase of the disease, not during the incubation^9^.

So far, the spread of COVID-19 has been exclusively contained by non-pharmacological health protection measures, especially social distancing, use of face masks, hand washing, isolation of confirmed cases, quarantine for close contacts and a number of constraints on several business activities as well as on social freedom of individuals^1,2^. Since these public health measures dramatically impact on the economy of countries and the personal life of people, they cannot be imposed for long time.

Therefore, there is a need to find pharmacological interventions to prevent and control COVID-19, supporting yet mitigating the existing IPC measures while an effective and safe vaccine will hopefully become available. Natural products derived from the innate immune system could be exploited as candidates for broad-spectrum antimicrobial agents. In this regard, hypothiocyanite (OSCN^−^) and lactoferrin (LF) can represent promising candidates for new anti-infective therapies.

OSCN^−^ is an anti-infectious agent, present in various exocrine secretions including the interface of the human central airways epithelium, that is produced by hydrogen peroxide (H_2_O_2_) and thiocyanate (anion SCN^−^) in presence of a peroxidase enzyme (lactoperoxidase, LPO). While it is present in the human bronchi^10^, LPO is nearly absent in the alveoli^11^. On the other hand, the angiotensin-converting enzyme 2 (ACE2), the receptor used by SARS-CoV-2 to enter cells^12^, is widely expressed in airways, particularly in nasal epithelial cells and type II alveolar epithelial cells^13^. The LPO-based system inactivates micro-organisms in the extracellular space, indeed the lack of LPO/H_2_O_2_/SCN^−^ in alveoli may result in devastating effects on ACE2-expressing alveolar type II cells^14^. In 2014, we have demonstrated for the first time that OSCN^−^ reduces the influenza virus A(H1N1)pdm09 infectivity when the virus is challenged with OSCN^−^ for 60 min at 37 °C before infection supporting the possibility to use it for influenza prevention and treatment^15^. Thereafter the antiviral activity of OSCN^−^ was tested also against several other strains of influenza virus, confirming a strain independent virucidal effect^16,17^.

LF is a natural multifunctional protein belonging to the family of transferrins, which can be found in human milk and external secretions as saliva, tears, milk, nasal and bronchial secretions, gastrointestinal fluids and urine mucosal secretions^18^. As a part of the innate human immune system, LF was found to be 150 fold over-expressed in SARS patients as compared to healthy controls^19^. LF has strong antiviral activity against a large range of DNA and RNA viruses^20^ and inhibits SARS-CoV-1 pseudovirus cell entry by binding the heparan sulfate proteoglycans of target cells^21^.

In view of the above considerations, we conducted an in-vitro study on OSCN^−^ and a combination of OSCN^−^ and LF, with the aim of testing their cytotoxic effect and virucidal activity against SARS-CoV-2.

## Virucidal activity of OSCN^−^ and LF

To investigate the virucidal activity of OSCN^−^ against SARS-CoV-2 we used a recombinant Vesicular Stomatitis Virus (rVSV), encoding the reported gene luciferase instead of the viral glycoprotein, pseudotyped with the Spike (S) protein of SARS-CoV-2 (rVSV-S). It has been shown that VSV can be pseudotyped by the S protein obtaining a virus with a tropism dictated by the heterologous S envelope which represents an excellent surrogate of SARS-CoV-2 to study virus entry and viral neutralization^22^. rVSV, as single replication step virus, can be manipulated under biosafety level 2 conditions, thereby overcoming the biosafety concern due to the manipulation of highly pathogenic viruses^23^. To this end, rVSV-S was incubated with different OSCN^−^ concentrations for 60 min at 37°C, then Vero cells were infected at multiplicity of infection (MOI) 0.065 focus forming unit (FFU) per cell. Sixteen hours post infection, cells were lysed and the luciferase expression was evaluated. As shown in Fig. 1a viral infection is inhibited in a dose dependent manner and the concentration capable of reducing the viral infectivity by 50% (IC50) was 4.64 μM, a value comparable with that we previously reported for influenza virus A(H1N1)pdm09^15^.

**Figure 1.**
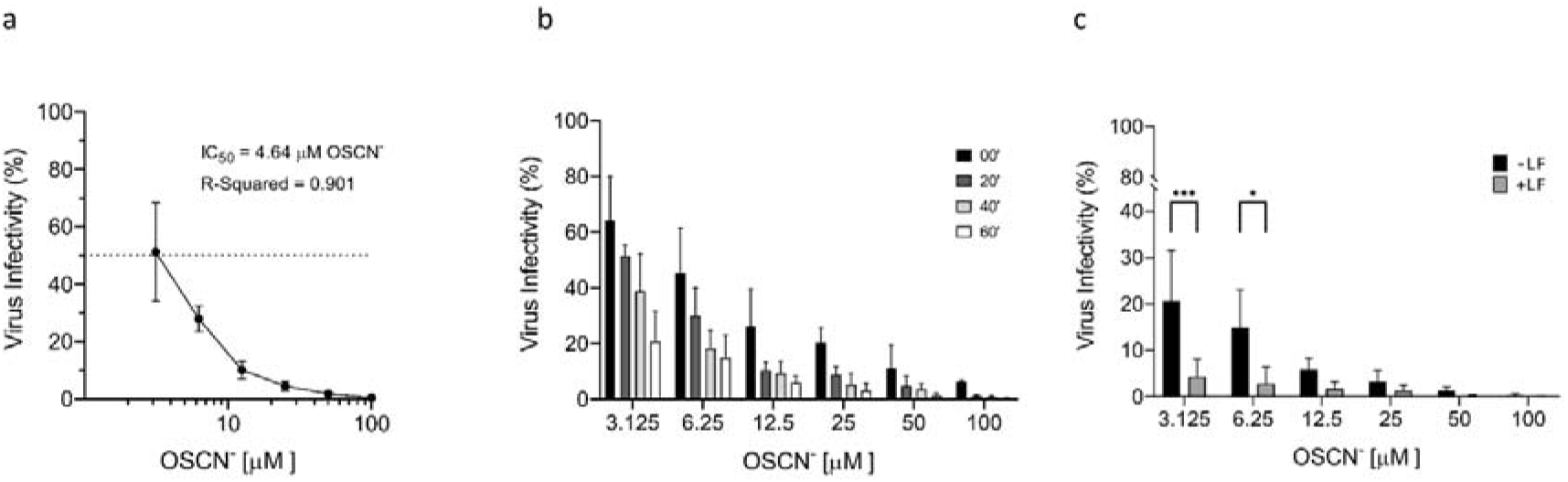
OSCN^−^ and OSCN^−^/LF virucidal activity against the pseudovirus VSV-S. **(a),** Efficiency of pseudovirus infection after exposition at different OSCN^−^ concentrations for 1 h at 37 °C. **(b)**, Evaluation of the virucidal activity of OSCN^−^ after pseudovirus treatment for 0, 20, 40, and 60 min at 37 °C before the infection of target cells. **(c)**, Comparison between OSCN^−^ and OSCN^−^+LF virucidal activity after 1 h of pretreatment of VSV-S before cells infection. Data (mean ± SD, N = 3, experiments in duplicate) are percentages of no drug, set as 100% (* = P < 0.05; *** = P < 0.001).

To evaluate the correlation between the time of virus exposure to OSCN^−^ and the virucidal activity, rVSV-S was incubated with OSCN^−^ for 60, 40 or 20 minutes before the infection of target cells (virus pre-treatment) or directly incubated (co-treatment) during the one hour of virus adhesion to cells (0 minutes, in Fig 1b). Although the efficiency of the virucidal activity was clearly time dependent in all the conditions tested, we observed a reduction of the viral infectivity of more than 50% until the concentration of 6.25 μM (Fig 1b), when the virus was incubated with OSCN^−^ before the infection. Moreover, starting from OSCN^−^ concentration of 12.5 μM a reduction of more than 50% was also observed for the virus treatment during the infection of cells. We also investigated the virucidal activity of OSCN^−^ and LF combination. Preliminary experiments with LF showed that concentrations of more than 1 g/L are required to inhibit viral infection (data not shown). Then, we selected a concentration of 4 g/L that was close to that previously used with OSCN^−^ in experiments against bacteria^24^. The combination OSCN^−^/LF showed a significant increase of the virucidal activity at lower doses of OSCN^−^, with an inhibition of viral infection of more than 90% in comparison to a reduction of 80-85% observed with OSCN^−^ (Fig. 1c) and 25% with LF alone (data not shown). These data suggest that, in our experimental conditions, OSCN^−^ has the main virucidal effect, while at lower OSCN^−^ concentrations LF can contribute to improve the virus inactivation. Of note, addition of OSCN^−^ and LF was not toxic to Vero cells, monitored for 24 h by the MTT test, indeed the inhibition of viral infectivity is due to the interference of the compounds on the capacity of the virus to infect cells (supplementary figure 1).

**Supplemental Figure 1.**
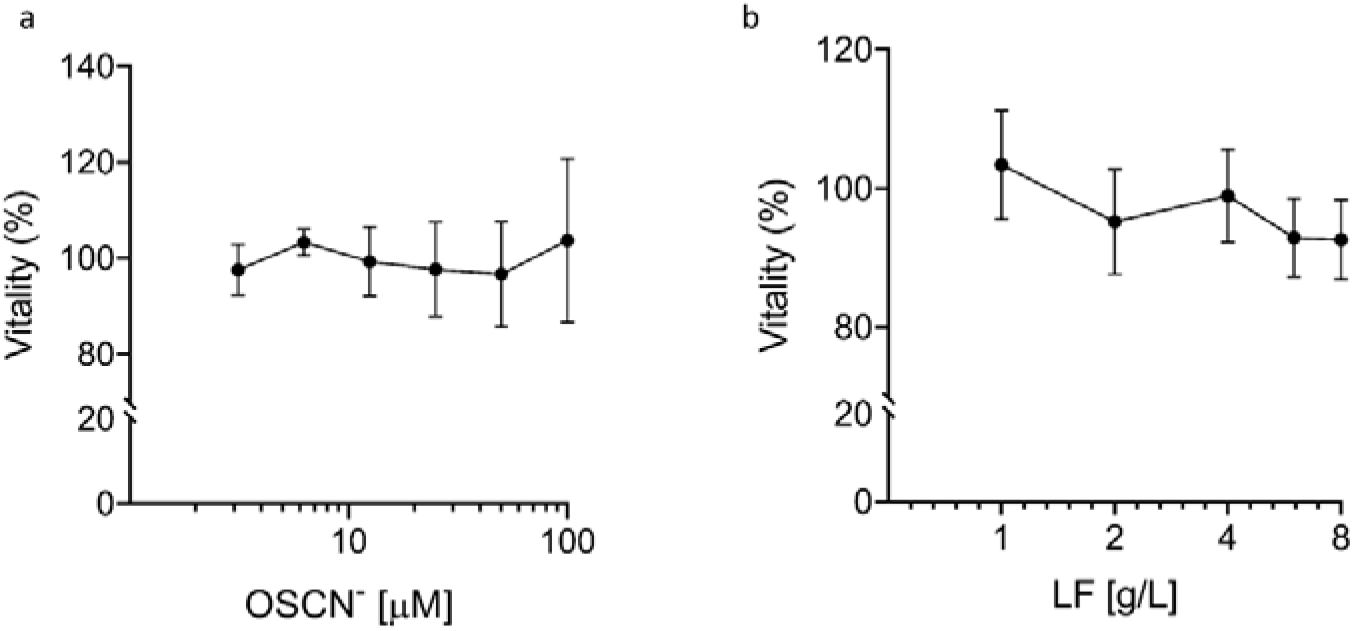
Cytotoxicity of OSCN^−^ and LF. The cytotoxicity of OSCN^−^ **(a)**and LF **(b)**was evaluated on Vero cells after 24 h of treatment using the MTT assay. Data (mean ± SD, N = 3, experiments in quadruplicate) are percentages of no drug, set as 100%.

Finally, to validate the results obtained with the rVSV-S, we performed experiments with OSCN^−^ and OSCN^−^/LF using SARS-CoV-2. After virus-compound incubation for 60 min, tenfold dilutions of virus-compound mix were inoculated on Vero-E6 cells (used for virus isolation and propagation) and the reduction of plaque generation was evaluated. Results reported in figure 2 confirmed the dose-dependent virucidal activity of OSCN^−^ (fig 2a) and demonstrated that higher doses of OSCN^−^ can efficiently inhibit SARS-CoV-2 infection also with an incubation of 20 min (figure 2b). In contrast to the pseudovirus, the enhancement of the virucidal activity of OSCN^−^ was less pronounced in presence of LF (figure 2c).

**Figure 2.**
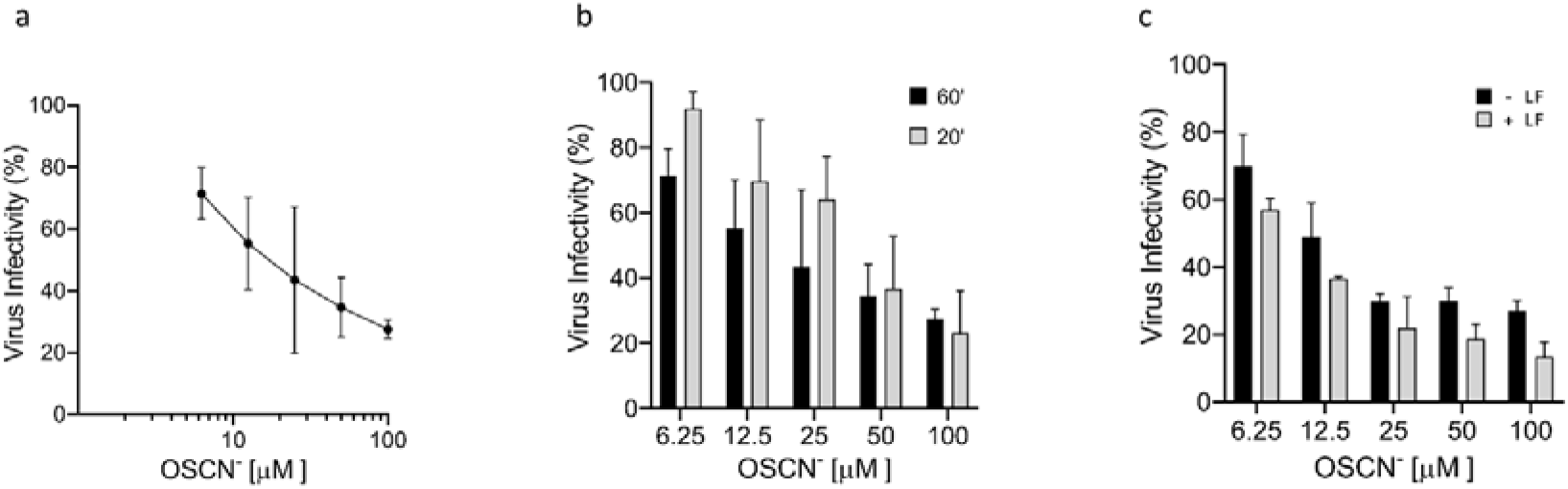
Virucidal activity of OSCN^−^ and OSCN^−^/LF against SARS-CoV-2. SARS-CoV-2 was treated for 1 h at 37 °C with OSCN^−^ alone or supplemented with LF before infection of cells. The reduction of infectivity was evaluated by plaque assay. **(a)**, virucidal effect of OSCN^−^. **(b)**, Comparison between different times of virus-OSCN^−^ exposition on the efficiency of the virucidal activity. **(c)**, Evaluation of the combination OSCN^−^+LF on the virucidal activity. Data (mean ± SD, N = 3, experiments in duplicate) are percentages of no drug, set as 100%.

Overall, our results indicate that OSCN^−^ has a strong virucidal activity against SARS-CoV-2, and the combination of OSCN^−^/LF further improved the OSCN^−^ virucidal effect by moderate margin.

## Conclusions

The pathogenesis of COVID-19 can be divided into three clinical stages depending on where the infection resides^14^. In the asymptomatic initial phase, COVID19 infection mostly resides in the nose where it elicits a minimal localized innate immune response. Scientists are looking for effective therapeutic strategies to reduce the viral shedding and disease spread. Very recently, a study evaluated the virucidal activity in-vitro of different commercially available oral rinses against SARS-CoV-2, assuming that oral rinsing might reduce the viral load of saliva and could thus lower the emission and transmission of SARS-CoV-2 through droplets. Viral infectivity of different SARS-CoV-2 strains was significantly reduced to undetectable levels by three formulations: Dequonal (active compounds: Dequalinium chloride and Benzalkonium chloride); Iso-Betadine mouthwash (Polyvidone-iodine) and Listerine Cool Mint (Ethanol and Essential oils)^25^. Thus, preliminary trials aimed to evaluate the reduction of viral shedding in confirmed COVID-19 patients have been registered^26,27^. In addition, it has been also shown that the mouth spray ColdZyme, used against common cold, inactivates SARS-CoV-2 in-vitro^28^. For the mild symptomatic phase, the infection is localized mostly in the pseudostratified epithelium of the larger airways and is accompanied by a more vigorous innate immune response. In the conducting airways, the epithelium can recover from the infection, because basal cells, progenitor for the bronchial epithelium, are spared. There may be more severe disease in the bronchioles, where club cells are likely infected. Mouthwashes and gargling are not effective treatments in this second phase, nor in the third (potentially lethal) phase of the disease, where the infection spreads into the gas exchange portion of the lung and infects ACE2-expressing alveolar type II cells. By contrast, given the absence of cytotoxic effect in our experimental conditions, OSCN^−^ and LF can enter the respiratory system in the form of aerosol and penetrate deep into the tracheobronchial tree up to alveolar sacs, inactivating both the viruses nearing the epithelium from outside and virions released from infected cells, thus mitigating or preventing all the three clinical stages of COVID-19 infection.

ALX-009, a combination of OSCN^−^ and bovine LF developed by ALAXIA SAS for the treatment of respiratory cystic fibrosis pathogens, has already being successfully tested in a phase 1 clinical trial (RCT02598999) on healthy volunteers and patients affected by cystic fibrosis^2^. Interestingly, OSCN^−^ and bovine LF concentrations in ALX-009 tested against pathogenic bacteria were considerably higher than those used in our experimental setting, reaching value of 3,600 μM and 8 g/L respectively, which therefore should likely maximize their respective virucidal activity^24^. Considering the current emergency situation, ALX-009 could be rapidly tried in clinical settings to verify its in-vivo efficacy, with the view of preventing the eventual progression of the disease in symptomatic and pre-symptomatic individuals testing positive for COVID-19 at nasal-pharyngeal swabs, as well as to reduce the virus shedding in asymptomatic subjects.

## Methods

### Cell culture and viruses

Vero (African green monkey kidney cells, ATCC^®^ CCL-81), Vero E6 (ATCC^®^ CRL1586) and HEK293T (ATCC^^®^^ CRL-11268TM) cells were grown in Dulbecco’s modified Eagle’s medium (D-MEM) containing 10% heat-inactivated fetal bovine serum (FBSi). Cells were maintained in a 5% CO_2_ incubator at 37□°C, routinely checked for mycoplasma and confirmed negative. Culture medium and FBSi were obtained from Gibco (Thermofisher).

SARS-CoV-2 (Genbank: MW000351) and the rVSVΔ;G-Luc, a recombinant Vesicular Stomatitis Virus containing the gene encoding for the luciferase protein in place of the VSV-G gene^29^, were used in the experiments.

### Virus stock preparation and titration

Vero E6 cells were seeded in T175 flasks and then infected with SARS-CoV-2 (MOI of ≈ 0.1). At 72-96 h post infection (p.i.), supernatants were collected, centrifuged at 2,300 rpm for 10 min, then stored in aliquots at −80 °C. Viral titer was determined by plaque assay on Vero E6 cells seeded on 24-well plates. Tenfold virus dilutions were prepared in DMEM and inoculated on confluent Vero-E6 cells for 1 h at 37 °C. After that, virus inoculum was removed from each well and cells were overlaid with 300 μL of 0.6% carboxymethylcellulose (Merck) diluted in DMEM supplemented with 2% FBSi. Seventy-two h p.i., cells were fixed adding 300 μL of 5% formaldehyde (Merck) in PBS 1x for 30 min at room temperature. Then, cells were stained with crystal violet in 20% ethanol. Virus titer was measured as plaque-forming units per milliliter (PFU/mL) based to the plaques formed in cell culture upon infection. All studies with viable SARS-CoV-2 were performed in the certified BSL3 laboratory of the Department of Molecular Medicine, University of Padova, in biological safety cabinets.

For SARS-CoV-2-pseudotyped VSV production (rVSV-S), HEK293T cells were seeded in T175 flasks and then transfected by calcium phosphate-DNA preciptation with 40 μg of pSARS-CoV-2-spike plasmid. After 24 h, cells were infected with the rVSVΔ;G-Luc virus at the multiplicity of infection (MOI) of 4 fluorescent focus-forming units (FFU)/cell. Sixteen hours p.i., cell culture supernatants were harvested and cell debris were cleared by centrifugation (2,300 rpm for 7 minutes at 4 °C). Then, virus particles were pelleted by ultracentrifugation on a 20 % p/v a sucrose cushion (27,000 rpm for 150 min at 4 °C) in a Beckmann SW 28 Ti Swinging-Bucket rotor. Pellets were resuspended in 1 mL of ice-cold PBS1X / tube and mixed. Subsequently, the virus was aliquoted and stored at −80 °C until use.

Titration of virus was determined by immunofluorescence on Vero cells seeded on 96-well plates. Viral stock was tenfold serially diluted in DMEM and inoculated on confluent Vero cells for 1 h at 37 °C. Then, cells were washed and DMEM supplemented with 10% FBSi was added. After 18 h, cells were fixed with precooled methanol-acetone for 1 h at −20 °C. Immune staining was performed by incubation with 1:3000 anti-VSV-M [23H12] monoclonal antibody (Kerafast) on the infected cells for 90 minutes at 37°C, followed by incubation with 1:1000 anti-rabbit Alexa Fluor^®^ 488 (Thermo Fisher Scientific) for 60 minutes at 37°C. The fluorescent foci in each well were counted and viral titer was expressed as focus-forming units per mL (FFU/mL)^30^.

### Compounds preparation and cytotoxicity assay

OSCN^−^ solution was prepared via enzymatic reaction with an automated equipment EOLEASE^®^ (Alaxia, Lyon, France). Due to its intrinsic reactivity, each solution freshly prepared was used alone or combined with Lactoferrin within 15 min after preparation. Reagents for the OSCN^−^ production were provided by Alaxia.

Pharma-grade lyophilized Lactoferrin (purity >98%) was made available by Alaxia, as sterile vials. Lyophilized Lactoferrin was reconstituted as solution with 0.9 % sodium chloride solution at 10 g/L.

Compound dilutions for virus treatment were performed in Minimum Essential Media (MEM) purchased by Gibco.

Cytotoxicity of OSCN^−^ and LF was determined on Vero and Vero-E6 cells after 24 h of treatment. Cell viability was tested with an assay based on the reduction of a tetrazolium salt (MTT Cell Proliferation Assay, Thermofisher) in a 96-well plate format according to the manufacturer’s instructions.

### Virucidal assay

To evaluate the OSCN^−^ and LF virucidal activity, 4 × 10^4^ FFU of rVSV-S were incubated for 1 h at 37 °C with 0 – 3.125 – 6.25 – 12.5 – 25 - 50 and 100 μM of OSCN^−^ with or without 4 g/L of LF. Positive control (mock sample) was treated with the solution used to prepare the OSCN^−^ simulating the max OSCN^−^ concentration. Next, Vero cells seeded on 96-well plates were infected for 1h at 37°C. Eighteen hours later, infection was evaluate measuring the relative light unit (RLU) with a VICTOR Multilabel Plate Reader (PerkinElmer) using the Steady-Glo^®^ Luciferase Assay System (Promega).

In the case of SARS-CoV-2, 1 × 10^5^ PFU were incubated for 1 h at 37 °C with 0 – 6.25 – 12.5 - 25 - 50 and 100 μM of OSCN^−^ with or without 4 g/L of LF. Next, tenfold virus dilutions were prepared in MEM and processed as above reported for the plaque assay. Viral titer was calculated for each sample and the virucidal activity was measured evaluating the efficiency of the infection in comparison to the mock treated control.

### Statistical analysis

All the experiments were performed in duplicate in at least three independent biological replicates. Statistical analysis was carried out with GraphPad Prism the one-way ANOVA test. The threshold for statistical significance was P<0.05. All details on sample size and P values for each experiment are provided in the relevant figure or its legend. Curve fitting was performed to determine IC50 values using a sigmoidal 4PL model in GraphPad Prism 8 software.

## Acknowledgements

We are grateful to Alaxia SAS that provided protocols, equipment and reagents to prepare OSCN^−^ and LF. We thank Michael Whitt, University of Tennessee, Memphis, USA, for providing the VSVΔ;G-Luc and Maria Rita Gismondo, L. Sacco University Hospital, Milan, Italy, for providing the SARS-CoV-2.

## Authors Contributions

L.C. and Cr.S. conceived the project and wrote the draft. Cr.S. propagate the SARS-CoV-2, performed experiments with SARS-CoV-2, provided reagents, and coordinated the project. Cl.S. performed experiments with SARS-CoV-2 and cell culture, collected and analyzed the data. M.M and M.V.S. performed experiments with rVSV-S and cell culture, collected and analyzed the data. G.M. contributed to the initial conceptualization of the project, and helped with data interpretation.

## Data availability statement

Raw data are available upon request.

## Additional information

Supplementary information is available for this paper

Correspondence and requests for materials should be addressed to L.C. or Cr.S.

